# Developing a modern data workflow for evolving data

**DOI:** 10.1101/344804

**Authors:** Glenda M. Yenni, Erica M. Christensen, Ellen K. Bledsoe, Sarah R. Supp, Renata M. Diaz, Ethan P. White, S.K. Morgan Ernest

## Abstract

Data management and publication are core components of the research process. An emerging challenge that has received limited attention in biology is managing, working with, and providing access to data under continual active collection. “Evolving data” present unique challenges in quality assurance and control, data publication, archiving, and reproducibility. We developed a evolving data workflow for a long-term ecological study that addresses many of the challenges associated with managing this type of data. We do this by leveraging existing tools to: 1) perform quality assurance and control; 2) import, restructure, version, and archive data; 3) rapidly publish new data in ways that ensure appropriate credit to all contributors; and 4) automate most steps in the data pipeline to reduce the time and effort required by researchers. The workflow uses two tools from software development, version control and continuous integration, to create a modern data management system that automates the pipeline.

## Introduction

Over the past few decades, biology has transitioned from a field where data are collected in hand-written notes by lone scientists, to an endeavor that increasingly involves large research teams coordinating data collection activities across multiple locations and data types. While there has been much discussion about the impact of this transition on the amount of data being collected (Hampton et al., 2013; Marx, 2013), there has also been a revolution in the frequency with which we collect those data. Instead of one-time data collection, biologists are increasingly asking questions and collecting data that require continually updating databases with new information. Long-term observational studies, experiments with repeated sampling, use of automatic sensors (e.g., temperature probes and satellite collars), and ongoing literature mining to build data compilations all produce continually-updating data. These data are being used to ask questions and design experiments that take advantage of regularly updating data streams: e.g., adaptive monitoring and management (Lindenmayer & Likens, 2009), iterative near-term forecasting (Dietze et al., 2018), detecting and preventing ecological transitions (Carpenter et al., 2011), and real-time cancer metabolism (Misun, Rothe, Schmid, Hierlemann, & Frey, 2016). Thus, whether studying changes in gene expression over time, or the long-term population dynamics of organisms, data that are being analyzed while they are still undergoing data collection is becoming a pervasive aspect of biology. This type of data has been referred to as “evolving data” (*sensu* Salzberg and Tsotras 1999, Ganti et al. 2001, Rauber et al. 2016; it is also referred to as “dynamic data”) to represent the idea that it is regularly changing and expanding.

Because evolving data are frequently updated, even during analysis, they present unique challenges for effective data management. These challenges have received little attention, especially regarding data that are collected by individual labs or small teams. All data must undergo quality assurance and quality control (QA/QC) protocols before being analyzed to find, correct, or flag inaccuracies due to data entry errors or instrument malfunctions. If data collection is finite, or if analysis will not be conducted until data collection is completed, these activities can be conducted on all of the data at once. Evolving data, however, are continually being collected, and new data require QA/QC before being added to the core database. This continual QA/QC demand places an extra burden on data managers and increases the potential for delays between when data are collected and when they are available to researchers to analyze. Thus, to be maximally useful, evolving datasets require protocols that promote rapid, ongoing data entry (either from field or lab notes or downloads from instrument data) while simultaneously detecting, flagging, and correcting data issues.

The need to analyze data still undergoing collection also presents challenges for managing data availability, both within research groups and while sharing with other research groups. By definition, continually-updating data regularly creates new versions of the data, resulting in different versions of the same dataset undergoing analysis at different times and by different researchers. Understanding differences in analyses over time or across researchers becomes more difficult if it is unclear which version of the data is being analyzed. This is particularly important for making research in biology more reproducible (Hampton et al., 2013; Errington et al., 2014). Efforts to share data with outside groups will encounter many of the same issues as sharing within a group. These challenges are magnified by the fact that the archiving solutions available to individual researchers (e.g. data papers, archiving of data as part of publications) treat data as largely static, which creates challenges for updating these data products. This static view of data publication also makes providing credit to data contributors challenging as new contributors become involved in collecting data for an existing data stream. Properly crediting data collectors is viewed as an essential component of encouraging the collection and open provision of valuable datasets (Reichman, Jones, & Schildhauer, 2011; Molloy, 2011). However, the most common approaches to citing and tracking data typically fail to properly acknowledge contributors to evolving datasets who join the project after the initial data paper or scientific paper is published, even when a more recent version is being analyzed.

Strategies for managing large amounts of continually-updated data exist in biology, but these are generally part of large, institutionalized data collection efforts with dedicated informatics groups, such as the U.S. National Ecological Observatory Network (NEON, https://www.neonscience.org), the National Center for Biotechnology Information (NCBI, https://www.ncbi.nlm.nih.gov), and the Australian Terrestrial Ecosystem Research Network (TERN, http://www.tern.org.au). As the frequency with which new data are added increases, it becomes more and more difficult for humans to manually provide data preparation and quality control (i.e., manual data checks, importing into spreadsheets for summarizing), making automated approaches increasingly important. Institutionalized data collection efforts include data management workflows to automate many aspects of the data management pipeline. These procedures include software that automates quality checks, flags data entry or measurement errors, integrates data from sensors, and adds quality-checked data into centralized database management systems. Developing systems like these typically requires dedicated informatics professionals, a level of technical support not generally available to individual researchers or small research groups that lack the funding and infrastructure to develop and maintain complex data management systems.

As a small group of researchers managing an ongoing, long-term research project, we have grappled with the challenges of managing evolving data and making them publicly available. Our research involves automated and manual data collection efforts, at daily through annual frequencies, conducted over forty years by a regularly changing group of personnel who all deserve credit for their contributions to the project. Thus, our experience covers much of the range of evolving data challenges that biologists are struggling to manage. We designed a modern workflow system to expedite the management of data streams ranging from weather data collected hourly by automated weather stations to plant and animal data recorded on datasheets in the field. We use a variety of tools that range from those commonly used in biology (e.g., MS Excel and programming in high-level languages like R or Python) to tools that biology is just beginning to incorporate (e.g., version control, continuous integration). Here, we describe the steps in our processes and the tools we use to allow others to implement similar evolving data systems.

## Implementing a modern data workflow

Setting up a data management system for automated management of continually-collected data may initially seem beyond the skill set of most empirically-focused lab groups. The approach we have designed, and describe below, does require some level of familiarity and comfort with computational tools such as a programming language (e.g., Python or R) and a version control system (e.g., git). However, data management and programming are increasingly becoming core skills in biology (Hampton et al., 2017), even for empirically-focused lab groups, and training in the tools we used to build an evolving data management system is available at many universities or through workshops at conferences. In designing and building the infrastructure for our study, our group consisted primarily of field ecologists, who received their training in this manner, and assistance from a computational ecologist for help figuring out overall design and implementation of some of the more advanced aspects. We have aimed this paper, and our associated tutorial, at empirical groups with little background in the tools or approaches we implemented. Our goal is to provide an introduction to the concepts and tools, general information on how such a system can be constructed, and assistance--through our tutorial--for building basic evolving data management systems. Readers interested in the specific details of our implementation are encouraged to peruse our active evolving data repository (www.github.com/weecology/PortalData).

## The model system

Our evolving data are generated by the Portal Project, a long-term study in ecology that is currently run by our research group (Ernest et al., 2018). The project was established in 1977 in the southwestern United States to study competition among rodents and ants, and the impact of these species on desert plants (Brown, 1998). This study produces several long-term evolving data sets. We collect these data at different frequencies (hourly, monthly, biannually, and annually), and each dataset presents its own challenges. Data on the rodents at the site are collected monthly on uniquely-tagged individuals. These data are the most time-intensive to manage because of how they are recorded (on paper datasheets), the frequency with which they are collected (every month), and the extra quality control efforts required to maintain accurate individual-level data. Data on plant abundances are collected twice a year on paper datasheets. These data are less intensive to manage because data entry and quality control activities are more concentrated in time and more limited in effort. We also collect weather data, generated hourly, which we download weekly from an automated weather station at the field site. Because we do not transcribe these data, there are no human-introduced errors. We perform weekly quality control efforts for these data, to check for issues with the sensors, including checking for abnormal values and comparing output to regional stations to identify extreme deviations from regional conditions. Given the variety of data that we collect, we require a generally flexible approach for managing the data coming from our study site. The diversity of evolving data that we manage makes it likely that our data workflow will address many of the data management situations that biologists collecting evolving data regularly encounter.

## Data Management Tools

To explain the workflow, we break it into steps focused on the challenges and solutions for each part of the overall data workflow (Figure 1). In the steps described below, we also discuss a series of tools we use which may not be broadly familiar across all fields of biology. We use R (R Development Core Team, 2018), an open-source programming language commonly used in ecology, to write code for acquiring and managing data and comparing files. We chose R because it is widely used in ecology and is a language our team was already familiar with. To provide a central place for storing and managing our data, we use GitHub (Box 1; https://github.com), an online service used in software development for managing version control. Version control systems are used in software development to provide a centralized way for multiple people to work on code and keep track of all the changes being made (Wilson et al., 2014). To help automate running our data workflow (so that it runs regularly without a person needing to manually run all the different pieces of code required for quality control, updating tables and other tasks), we expand on the idea of continuous analysis proposed by Beaulieu-Jones and Greene (2017) by using a continuous integration service to automate data management (see Box 2). In a continuous integration workflow, the user designates a set of commands (in our case, this includes R code to error-check new data and update tables), which the continuous integration service runs automatically when data or code is updated or at user-specified times. We use a continuous integration service called Travis (https://travis-ci.com/), but there are several other options available including other services (e.g., AppVeyor https://www.appveyor.com/) and systems that can be run locally (e.g., Jenkins; https://jenkins.io/). Other tools are used for only small, distinct tasks in the pipeline and are described as needed. All of the code we use in our data management process can be found in our GitHub repository (https://github.com/weecology/PortalData) and is archived on Zenodo (https://zenodo.org/record/1219752).

**Figure 1:**
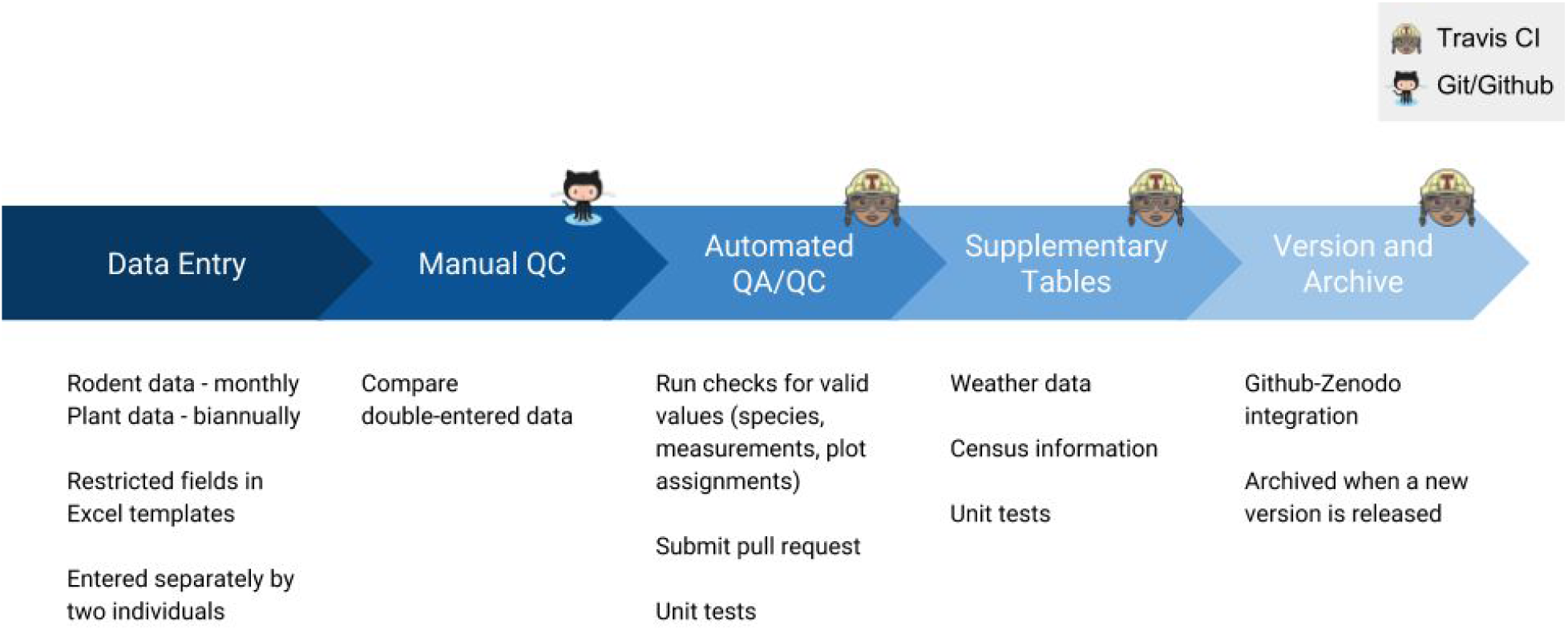
Our data workflow

### QA in data entry

For data collected onto datasheets in the field, the initial processing requires human interaction to enter the data and check that data entry for errors. Upon returning from the field, new data are manually entered into Excel spreadsheets by two different people. We use the “data validation” feature in Excel to restrict possible entries as an initial method of quality control. This feature is used to restrict accepted species codes to those on a pre-specified list and restrict the numeric values to allowable ranges. The two separately-entered versions are compared to each other using an R script to find errors from data entry. The R script detects any discrepancies between the two versions and returns a list of row numbers in the spreadsheet where these discrepancies occur, which the researcher then uses to compare to the original data sheets and fix the errors.

### Adding data to databases on GitHub

To add data (or correct errors) to our master copy of the database, we use a system designed for managing and tracking changes to files called version control. Version control was originally designed for tracking changes to software code, but can also be used to track changes to any digital file, including datafiles (Ram 2013, Pröll and Meixner 2016). We use a specific version control system, git and the associated GitHub website, for managing version control (see Box 1 for details; https://www.github.com). We store the master version of the Portal data files on GitHub’s website (https://github.com/weecology/PortalData). The data, along with the code for data management, are stored in the version control equivalent of a folder, called a repository. Through this online repository, everyone in the project has access to the most up-to-date, or “master”, version of both the data and the data management code. To add or change data in this central repository we edit a copy of the repository on a user’s local computer, save the changes along with a message describing their purpose, and then send a request through GitHub to have these changes integrated into the central repository (Box 1). This version control based process retains records of every change made to the data along with an explanation of that change (Ram 2013, Pröll and Meixner 2016). It also makes it possible to identify changes between different stages and go back to any previous state of the data. As such, it protects data from accidental changes and makes it easier to understand the provenance of the data.

### Automated QA/QC and human review

Another advantage of this version control based system is that it makes it relatively easy to automate QA/QC checks of the data and facilitates human review of data updates. Once the researcher has updated their local copy of the database they create a “pull request” (i.e. a request for someone to pull the user’s changes into the master copy). This request automatically triggers the continuous integration system to run a predetermined set of QA/QC checks. These QA/QC checks check for validity and consistency both within the new data (e.g., checking that all plot numbers are valid and that every quadrat in each plot has data recorded) and between the old and new data (e.g., ensuring that species identification is consistent for recaptured rodents with the same identifying tag). This QA/QC system is essentially a series of unit tests on the data. Unit testing is a software testing approach that checks to make sure that pieces of code work in the expected way (Wilson et al., 2014). We use tests, written using the ‘testthat’ package (Wickham, 2011), to ensure that all data contain consistent, valid values. If these checks identify issues with the data they are automatically flagged in the pull request indicating that they need to be fixed before the data are added to the main repository. The researcher then identifies the proper fix for the issue, fixes it in their local copy, and updates the pull request, which is then retested to ensure that the data pass QA/QC before it is merged into the main repository.

In addition to automated QA/QC, we also perform a human review of any field entered data being added to the repository. At least one other researcher--specifically *not* the researcher who initiated the pull request--reviews the proposed changes to identify any potential issues that are difficult to identify programmatically. This is facilitated by the pull request functionality on GitHub, which shows this reviewer only the lines of data that have have been changed. Once the changes have passed both the automated tests and human review, a user confirms the merge and the changes are incorporated into the master version of the database. Records of pull requests that have been merged with the main dataset are retained in git and on GitHub, and it is possible to revert to previous states of the data at any time.

### Automated updating of supplemental tables

Once data from the field is merged into the main repository, there are several supplemental data tables that need to be updated. These supplemental tables often contain information about each data collection event (e.g., sampling intensity, timing) that cannot be efficiently stored in the main data file. For example, as a supplemental table to our plant quadrat data, we have a separate table containing information on whether or not each of the 384 permanent quadrats was sampled during each sampling period. This table allows us to distinguish “true zeros” from missing data. Since this information can be derived from the entered data, we have automated the process of updating this table (and others like it) in order to reduce the time and effort required to incorporate new sampling events into the database. For each table that needs to be updated, we wrote a function to: i) confirm that the supplemental table needs to be updated, ii) extract the relevant information from the new data in the main data table, and iii) append the new information to the supplemental table. The update process is triggered by the addition of new data into one of the main data tables, at which point the continuous integration service executes these functions (see Box 2). As with the main data, automated unit tests ensure that all data values are valid and that the new data are being appended correctly. Automating curation of these supplemental tables reduces the potential for data entry errors and allows researchers to allocate their time and effort to tasks that require intellectual input.

### Automatically integrating data from sensors

We collect weather data at the site from an on-site weather station that transmits data over a cellular connection. We also download data from multiple weather stations in the region whose data is streamed online. We use these data for ecological forecasting (White et al., 2018) which requires the data to be updated in the main database in near real-time. While data collected by automated sensors do not require steps to correct human-entry errors, they still require QA/QC for sensor errors and the raw data need to be processed into the most appropriate form for our database. To automate this process, we developed R scripts to download the data, transform them into the appropriate format, and automatically update the weather table in the main repository. This process is very similar to that used to automatically update supplemental tables for the human-generated data. The main difference is that, instead of humans adding new data through pull requests, we have scheduled the continuous integration system to download and add new weather data weekly. Since weather stations can produce erroneous data due to sensor issues (our station is occasionally struck by lightning resulting in invalid values), we also run basic QA/QC checks on the downloaded data to make sure the weather station is producing reasonable values before the data are added. Errors identified by these checks will cause our continuous integration system to register an error indicating that they need to be fixed before the data will be added to the main repository (similar to the QA/QC process described above). This process yields fully automated collection of weather data in near real-time. Automation of this process has the added benefit of allowing us to monitor conditions in the field and the weather station itself. We know what conditions are like at the site in advance of trips to the field and if there are issues with the weather station we can come prepared to fix them rather than discovering the problem unexpectedly when we arrive at our remote field site.

### Versioning

A common issue with evolving datasets is that the data available at one point in time are not the same as the data at some point in the future. The evolving nature of evolving data can cause difficulties for precisely reproducing prior analyses. This issue is rarely addressed, and when it is the typical approach is only noting the date on which the data were accessed. Noting the date acknowledges the continually changing state of the data but does not address reproducibility issues unless copies of the data for every possible access date are available. In order to support reproducibility of analyses on evolving data, Rauber et al. (2016) suggest the use of data versioning and timestamping of changes so that analyses can be rerun on a specific. To address this issue, we automatically make a “release” every time new data are added to the database using the GitHub API. This is modeled on the concept of releases in software development, where each “release” points to a specific version of the software that can be accessed and used in the future even as the software continues to change. By giving each change to the data a unique release code (known as a “version”), the specific version of the data used for an analysis can be referenced directly, and this exact form of the data can be downloaded to allow fully reproducible analyses even as the dataset is continually updated. This solves a commonly experienced reproducibility issue, that occurs both within and between labs, where it is unclear whether differences in results are due to differences in the data or the implementation of the analysis. We name the versions following the newly developed Frictionless Data data-versioning guidelines, where data versions are composed of three numbers: a major version, a minor version, and a “patch” version (https://frictionlessdata.io/specs/patterns/). For example, the current version of the datasets is 1.34.0, indicating that the major version is 1, the minor version is 34, and the patch version is 0. The major version is updated if the structure of the data is changed in a way that would break existing analysis code. The minor version is updated when new data are added, and the patch version is updated for fixes to existing data.

### Archiving

Through GitHub, researchers can make their data publicly available by making the repository public, or they can restrict access by making the repository private and giving permissions to select users. While repository settings allow data to be made available within or across research groups, GitHub does not guarantee the long-term availability of the data. GitHub repositories can be deleted at any time by the repository owners, resulting in data suddenly becoming unavailable (Bergman, 2012; White, 2015). To ensure that data are available in the long-term (and satisfy journal and funding agency archiving requirements), data also need to be archived in a location that ensures data availability is maintained over long periods of time (Bergman, 2012; White, 2015). While there are a variety of archiving platforms available (e.g., Dryad, FigShare), we chose to permanently archive our data on Zenodo, a widely used general purpose repository that is actively supported by the European Commission. We chose Zenodo because there is already a GitHub integration that automatically archives the data to Zenodo every time a release is made in the GitHub repository. In some fields, disciplinary repositories are the best choice for archiving some kinds of data. To make this feasible for evolving data, these repositories will need to support automatic updating (e.g., via an API) and data versioning. Currently the disciplinary repositories for ecological data (i.e., Dryad and Ecological Archives), and many other disciplines, lack these capabilities. We recommend that data archives follow the lead of *Earth System Science Data* by incorporating an explicit focus on evolving data (or “living data” as they refer to it; https://www.earth-system-science-data.net/living_data_process.html).

Zenodo incorporates the versioning described above so that version information is available in the permanently archived form of the data. Each version receives a unique DOI (Digital Object Identifier) to provide a stable web address to access that version and a top-level DOI is assigned to the entire archive, which can be used to collectively reference all versions of the dataset. This allows someone publishing a paper using the Portal Project data to cite the exact version of the data used in their analyses to allow for fully reproducible analyses and to cite the dataset as a whole to allow accurate tracking of the usage of the dataset.

### Citation and authorship

Providing academic credit for collecting and sharing data is essential for a healthy ecosystem supporting data collection and reuse (Reichman, Jones, & Schildhauer, 2011; Molloy, 2011). The traditional solution has been to publish “data papers” that allow a dataset to be treated like a publication for both reporting as academic output and tracking impact and usage through citation. This is how the Portal Project has been making its data openly available for the past decade, with data papers published in 2009 and 2016 (Ernest et al., 2009; Ernest et al., 2016). Because data papers are modelled after scientific papers they are static in nature and therefore have two major limitations for use with evolving data. First, the current publication structure does not lend itself to data that are regularly updated. Data papers are typically time-consuming to put together, and there is no established system for updating them. The few long-term studies that publish data papers have addressed this issue by publishing new papers with updated data roughly once every five years (e.g., Ernest et al. 2009 and 2016, Clark and Clark, 2000 and 2006). This does not reflect that the dataset is a single growing entity and leads to very slow releases of data. Second, there is no mechanism for updating authorship on a data paper as new contributors become involved in the project. In our case, a new research assistant joins the project every one to two years and begins making active contributions to the dataset. Crediting these new data collectors requires updating the author list while retaining the ability of citation tracking systems like Google Scholar to track citation. An ideal solution would be a data paper that can be updated to include new authors, mention new techniques, and link directly to continually-updating data in a research repository. This would allow the content and authorship to remain up to date while recognizing that the dataset is a single living entity. We have addressed this problem by writing a data paper (Ernest et al., 2018) that currently resides on bioRxiv, a pre-print server widely used in the biological sciences. BioRxiv allows us to update the data paper with new versions as needed, providing the flexibility to add information on existing data, add new data that we have made available, and add new authors. Like the Zenodo archive, BioRxiv supports versioning of preprints, which provides a record of how and when changes were made to the data paper and authors are added. Google Scholar tracks citations of preprints on bioRxiv, providing the documentation of use that is key to tracking the impact of the dataset and justifying its continued collection to funders.

### Open licenses

Open licenses can be assigned to public repositories on GitHub, providing clarity on how the data and code in the repository can be used (Wilson et al., 2014). We chose a CC0 license that releases our data and code into the public domain, but there are a variety of license options that users can assign to their repository specifying an array of different restrictions and conditions for use. This same license is also applied to the Zenodo archive.

## Discussion

Data management and sharing are receiving increasing attention in science, resulting in new requirements from journals and funding agencies. Discussions about modern data management focus primarily on two main challenges: making data used in scientific papers available in useful formats to increase transparency and reproducibility (Reichman, Jones, & Schildhauer, 2011; Molloy, 2011) and the difficulties of working with exceptionally large data (Marx, 2013). An emerging data management challenge that has received significantly less attention in biology is managing, working with, and providing access to data that are undergoing continual active collection. These data present unique challenges in quality assurance and control, data publication, archiving, and reproducibility. The workflow we developed for our long-term study, the Portal Project (Ernest et al., 2018), solves many of the challenges of managing this “evolving data”. We employ a combination of existing tools to reduce data errors, import and restructure data, archive and version the data, and automate most steps in the data pipeline to reduce the time and effort required by researchers. This workflow expands the idea of continuous analysis (*sensu* Beaulieu-Jones and Greene, 2017) to create a modern data management system that uses tools from software development to automate the data collection, processing, and publication pipeline.

We use our evolving data management system to manage data collected both in the field by hand and automatically by machines, but our system is applicable to other types of data collection as well. For example, teams of scientists are increasingly interested in consolidating information scattered across publications and other sources into centralized databases: e.g., plant traits (Kattge et al., 2011), tropical diseases (Hürlimann et al., 2011), biodiversity time series (Dornelas & Willis, 2017), vertebrate endocrine levels (Vitousek et al., 2018), and microRNA target interactions (Chou et al., 2016). Because new data are always being generated and published, literature compilations also have the potential to produce evolving data like field and lab research. Whether part of a large, international team such as the above efforts or single researchers interested in conducting meta-analyses, phylogenetic analyses, or compiling DNA reference libraries for barcodes, our approach is flexible enough to apply to most types of data collection activities where data need to be ready for analysis before the endpoint is reached.

The main limitation on the infrastructure we have designed is that it cannot handle truly large data. Online services like GitHub and Travis typically limit the amount of storage and compute time that can be used by a single project. GitHub limits repository size to 1 GB and file size to 100 MB. As a result, remote sensing images, genomes, and other data types requiring large amounts of storage will not be suitable for the GitHub-centered approach outlined here. Travis limits the amount of time that code can run on its infrastructure for free to one hour. Most research data and data processing will fit comfortably within these limits (the largest file in the Portal database is currently <20 MB and it takes <15 minutes for all data checking and processing code to run), so we think this type of system will work for the majority of research projects. However, in cases where larger data files or longer run times are necessary, it is possible to adapt our general approach by using equivalent tools that can be run on local computing resources (e.g, GitLab for managing git repositories and Jenkins for continuous integration) and using tools that are designed for versioning large data (e.g., dat; Ogden, McKelvey, & Madsen, 2017; or git Large File Storage; Perez-Riverol et al. 2016).

One advantage of our approach to these challenges is that it can be accomplished by a small team composed of primarily empirical researchers. However, while it does not require dedicated IT staff, it does require some level of familiarity with tools that are not commonly used in biology. To implement this approach, many research groups will need computational training or assistance. The use of programming languages for data manipulation, whether in R, Python, or another language, is increasingly common, and many universities offer courses that teach the fundamentals of data science and data management (e.g., http://www.datacarpentry.org/semester-biology/). Training activities can also be found at many scientific society meetings and through workshops run by groups like The Carpentries, a non-profit group focused on teaching data management and software skills--including git and GitHub--to scientists (https://carpentries.org/). A set of resources for learning the core skills and tools discussed in this paper is provided in Box 3. The most difficult to learn tool is continuous integration, both because it is a more advanced computational skill not covered in most biology training courses, and because existing documentation is primarily aimed at people with high levels of technical training (e.g., software developers). To help researchers implement this aspect of the workflow, including the automated releasing and archiving of data, we have created a starter repository including reusable code and a tutorial to help researchers set up continuous integration and automated archiving using Travis for their own repository (http://github.com/weecology/livedat). The value of the tools used here emphasizes the need for more computational training for scientists at all career stages, a widely recognized need in biology (Barone, Williams, & Micklos, 2017; Hampton et al., 2017). Given the importance of rapidly available evolving data for forecasting and other research, training, supporting, and retaining scientists with advanced computational skills to assist with setting up and managing evolving data workflows will be an increasing need for the field.

Evolving data is a relatively new data type for biology and one that comes with a unique set of computational challenges. While our data management approach provides a prototype for how research groups without dedicated IT support can construct their own pipelines for managing this type of data, continued investment in this area is needed. Our hope is that our approach serves as a catalyst for tool development that makes implementing evolving data management protocols increasingly accessible. Investments in this area could include improvements in tools implementing continuous integration, performing automated data checking and cleaning, and managing evolving data. Additional training in automation and continuous analysis for biologists will also be important for helping the scientific community advance this new area of data management. These investments will help decrease the current management burden of evolving data, which will allow researchers to make data available more quickly and effectively and let them spend more time collecting and analyzing data than managing it.

## Acknowledgements

This research, E. Christensen, and E. Bledsoe were all supported by the National Science Foundation through grant 1622425 to S.K.M. Ernest and by the Gordon and Betty Moore Foundation’s Data-Driven Discovery Initiative through grant GBMF4563 to E.P. White. R.M. Diaz was supported by a National Science Foundation Graduate Research Fellowship (DGE-1315138).

## Boxes

### Box 1: Version controlling data using git and Github

Version control systems are a set of tools for continually tracking and archiving changes made to a set of files. These systems were originally designed to facilitate collaborative work on software that was being continuously updated but can also be used when working with moderately sized data files. Version control tracks information about changes to files using “commits,” which record the precise changes made to a file or group of files along with a message describing why those changes were made. We use one of the most popular version control systems, git, along with an online system for managing shared git repositories, GitHub.

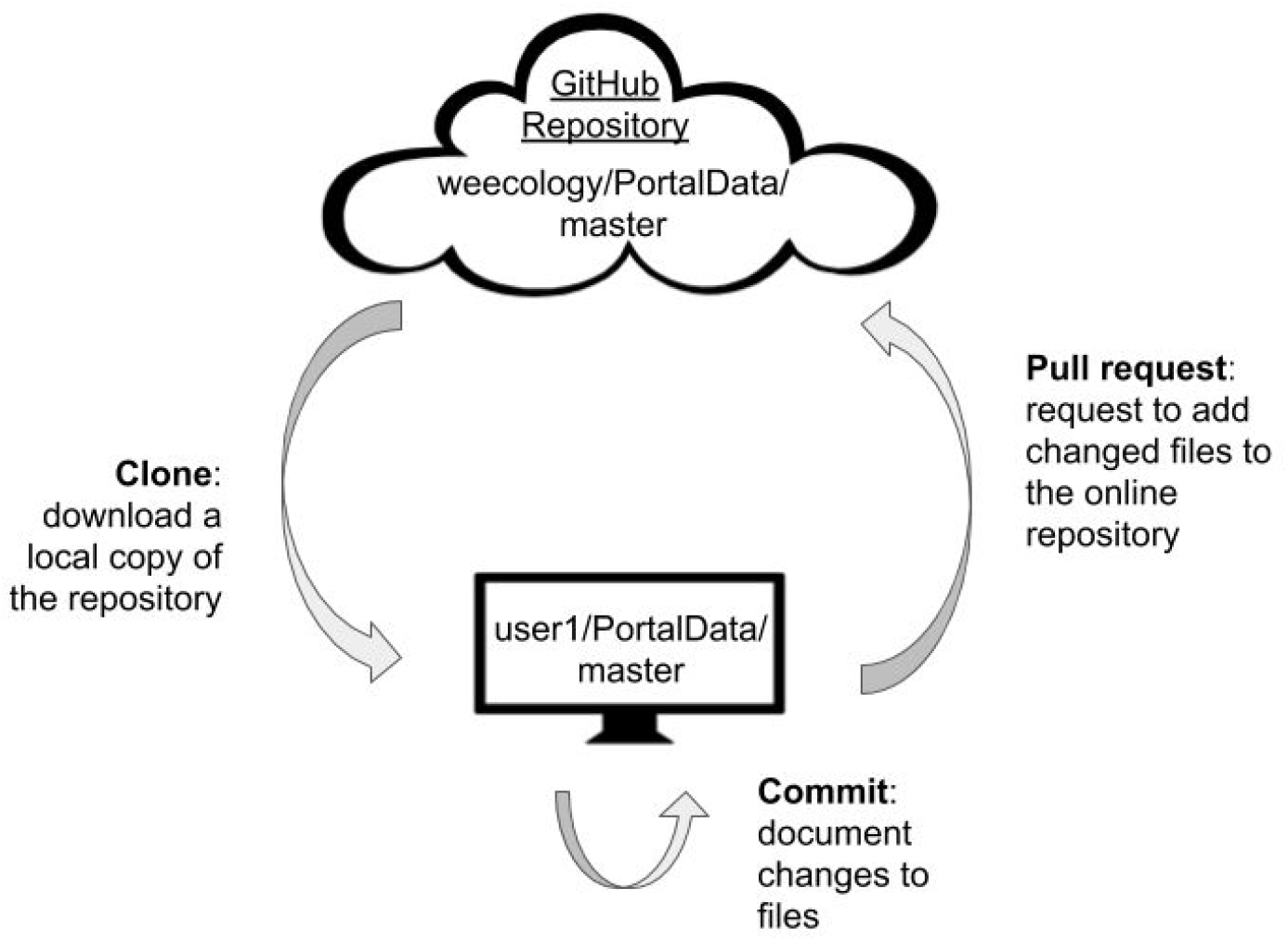

Version controlled projects are stored in “repositories,” (akin to a folder) and there is typically a central copy of the repository online to allow collaboration. In our case, this is our main GitHub repository that is considered to be the official version of the data (https://github.com/weecology/PortalData). Users can edit this central repository directly, but usually users create their own copies of the main repository called “forks” or “clones”. Changes made to these copies do not automatically change the main copy of the repository. This allows users to have one or more copies of the master version where they can make and check changes (e.g., adding data, changing data-cleaning code) before they are added to the main repository. As the user makes changes to their copy of the repository, they document their work by “committing” their changes. The version control system maintains a record of each commit, and it is possible to revert to past states of the data at any time. Once a set of changes is complete, they can be “merged” into the main repository through a process called a “pull request”. A pull request is a request by a user for someone to merge their changes into the main repository holding the primary copy of the data or code (a request that your changes be “pulled” into the main repository). As part of the pull request process, Github highlights all of the changes from the master version (additions or deletions), making it easy to see what changes are being proposed and determine whether they are good changes to make. Pull requests can also be automatically tested to make sure that the proposed changes do not alter the core functionality of the code or the core requirements of the data. Once the pull request is accepted, those changes become part of the main repository, but can be undone at any time if needed.

### Box 2: Travis

Continuous integration is a practice used in software engineering to automate testing and integrate new code into the main code base of a project. While designed as a software development tool, continuous integration has features which are useful for automating the management of evolving data: it detects changes in files, automates running code, and tests output for consistency. Because these tasks are also useful in a research context, this lead to the suggestion that continuous analysis could be used to drive research pipelines (Beaulieu-Jones and Greene, 2017). We expand on this concept by applying continuous integration to the management of evolving data.

The continuous integration service that we use to manage our evolving data is Travis (travis-ci.org), which integrates easily with Github. We tell Travis which tasks to perform by including a .travis.yml file (example below) in the GitHub repository containing our data, which is then executed whenever Travis is triggered.

Below is the Portal Data .travis.yml file and how it specifies the tasks Travis is to perform. First, Travis runs an R script that will install all R packages listed in the script (the “install:” step). It then executes a series of R scripts that update tables and run QA/QC tests in the Portal Data repository (the “script:” step):

- Update the regional weather tables [line 10]
- Run the tests (using the testthat package) [line 11]
- Update the weather tables from our weather station [line 12]
- Update the rodent trapping table (if new rodent data have been added, this table will grow, otherwise it will stay the same) [line 13]
- Update the plots table (if new rodent data have been added, this table will grow, otherwise it will stay the same) [line 14]
- Update the new moons table (if new rodent data have been added, this table will grow, otherwise it will stay the same) [line 15]
- Update the plant census table (if new plant data have been added, this table will grow, otherwise it will stay the same) [line 16]

If any of the above scripts fail, the build will stop and return an error that will help users determine what is causing the failure.

Once all the above steps have successfully completed, Travis will perform a final series of tasks (the “after_success:” step):

1. Make sure Travis’ session is on the master branch of the repo
2. Run an R script to update the version of the data (see the versioning section for more details)
3. Run a script that contains git commands to commit new changes to the master branch of the repository.

~~~
.travis.yml:
1 language: r
2 cache: packages
3 sudo: false
4 warnings_are_errors: false
5
6 install:
7    - Rscript install-packages.R
8
9 script:
10    - R -e ′setwdC (″DataCleaningScripts″); source(″get_regional_weather.R″); append_regional_weather()′
11    - Rscript testthat.R
12    - R -e ′setwdC (″DataCleaningScripts″); source(″new_weather_data.R″); append_weather()′
13    - R -e ′setwdC (″DataCleaningScripts″); source(″update_portal_rodent_trapping.r″); writetrappingtable()′
14    - R-e ′setwdC (″DataCleaningScripts″); source(″update_portal_plots.R″); writeportalplotsO ′
15    - R -e ′setwdC (″DataCleaningScripts″); source(″new_moon_numbers. r″); writenewmoons()′
16     - R -e ′setwdC (″DataCleaningScripts″); source(″update_portal_plant_censuses.R″); writecensustable() ′
17
18 after_success:
19    - git checkout master
20    - Rscript update_version.R
21    - bash update_repo.sh
~~~

Travis not only runs on the main repository, but also runs its tests on pull requests before they are merged. This automates the QA/QC and allows detecting data issues before changes are made to the main datasets or code. If the pull request causes no errors when Travis runs it, it is ready for human review and merging with the repository. After merging, Travis runs again in the master branch, committing any changes to the data to the main database. Travis runs whenever pull requests are made or changes detected in the repository, but can also be scheduled to run automatically at time intervals specified by the user, a feature we use to download data from our automated weather station.

### Box 3: Resources

#### Get Started

Evolving data Starter Repository: http://github.com/weecology/livedat

Open Source Licenses: https://choosealicense.com/

Unit Testing with the testthat package: http://r-pkgs.had.co.nz/tests.html

Data Validation in Excel: https://support.microsoft.com/en-us/help/211485/description-and-examples-of-data-validation-in-excel

Stack Overflow: https://stackoverflow.com/

#### Git/Git Hosts

Resources to learn git: https://try.github.io/

GitHub Learning Lab: https://lab.github.com/

Learn Git with Bitbucket: https://www.atlassian.com/git/tutorials/learn-git-with-bitbucket-cloud

Get Started with GitLab: https://docs.gitlab.com/ee/intro/

GitHub-Zenodo Integration: https://guides.github.com/activities/citable-code/

#### Continuous Integration

Version Control for Beginners: https://www.atlassian.com/git/tutorials

Travis Core Concepts for Beginners: https://docs.travis-ci.com/user/for-beginners/

Getting Started with Travis: https://docs.travis-ci.com/user/getting-started/

Getting Started with Jenkins: https://jenkins.io/doc/pipeline/tour/getting-started/

Jenkins learning resources: https://dzone.com/articles/the-ultimate-jenkins-ci-resources-guide

#### Training

The Carpentries: https://carpentries.org/

Data Carpentry: http://www.datacarpentry.org/

Software Carpentry: https://software-carpentry.org/

### Glossary

**CI/continuous integration**: (also see Box 2) the continuous application of quality control. A practice used in software engineering to continuously implement processes for automated testing and integration of new code into a project.

**Git**: (also see Box 1) Git is an open source program for tracking changes in text files (version control), and is the core technology that GitHub, the social and user interface, is built on top of.

**GitHub**: (also see Box 1) a web-based hosting service for version control using git.

**Github-Travis integration**: connects the Travis continuous integration service to build and test projects hosted at GitHub. Once set up, a GitHub project will automatically deploy CI and test pull requests through Travis.

**Github-Zenodo integration**: connects a Github project to a Zenodo archive. Zenodo takes an archive of your GitHub repository each time you create a new release.

**Evolving data**: data that continue to be updated and added to, while simultaneously being made available for analyses. For example: long-term observational studies, experiments with repeated sampling, data derived from automated sensors (e.g., weather stations or GPS collars).

**Pull request**: A set of proposed changes to the files in a GitHub repository made by one collaborator, to be reviewed by other collaborators before being accepted or rejected.

**QA/QC**: Quality Assurance/Quality Control. The process of ensuring the data in our repository meet a certain quality standard.

**Repository**: a location (folder) containing all the files for a particular project. Files could include code, data files, or documentation. Each file’s revision history is also stored in the repository.

**testthat**: an R package that facilitates formal, automated testing

**Travis CI**: (also see Box 2) a hosted continuous integration service that is used to test and build GitHub projects. Open source projects are tested at no charge.

**unit test**: a software testing approach that checks to make sure that pieces of code work in the expected way

**Version control**: A system for managing changes made to a file or set of files over time that allows the user to a) see what changes were made when and b) revert back to a previous state if desired

**Zenodo**: a general, open-access, research data repository

